# Beekeepers’ preferences for honeybee breeding goals: a French case study

**DOI:** 10.64898/2026.01.28.701934

**Authors:** T. Kistler, B. Basso, A. Lauvie, F. Phocas

**Author notes:** Corresponding author: Tristan Kistler. Co-first authors: Tristan Kistler, Benjamin Basso.

## Abstract

Honeybee breeding plans are relatively recent in most countries. In France, diverse small-scale breeding groups are emerging. Beekeepers are highly diverse in their motivations, farm productions and services, practices and management techniques. Yet, little is known about what beekeepers would consider as relevant breeding goals in the design of breeding plans. We therefore conducted an online survey answered by about 250 French beekeepers, mostly professionals, to assess their perceived importance of including 20 pre-defined traits in breeding goals and to identify how beekeeping profiles might influence these priorities. Respondents rated each trait as essential, useful, or useless, and indicated if they wished useful or essential traits to be genetically improved or merely maintained at their current level. Results indicated a strong preference for multi-trait selection, with a median of 13 traits considered useful or essential. Honey yield, disease resistance, swarming tendency, gentleness, and summer feed autonomy, emerged as the main traits of interest with about 90% of beekeepers finding them at least useful. About 40% or more only wished to maintain these traits at their current level rather than to directionally improve them. A major exception to this was disease resistance, that 75% wanted to improve. Bees’ genetic background influenced the most the importance attributed to breeding goal traits, while other beekeeping profile characteristics only had a marginal effect on breeding goal trait priorities.

Some poorly studied traits, such as summer and winter feed autonomy, winter diapause, and longevity, were considered at least useful in a breeding goal by over 70% of beekeepers. Future research is needed to explore possible selection criteria for these traits and estimate the potential for their genetic improvement.

**Implications:** Our survey shows that French beekeepers wish to improve or maintain through selective breeding usual colony production and behavioral traits, but also colony resilience, especially disease resistance and feed autonomy. However, trait priorities differ depending on the genetic background of the bees used. This knowledge is essential for designing breeding programs that truly match beekeeper needs and for identifying which traits deserve research attention. In France, beekeepers are increasingly starting breeding efforts to adapt their bees to current conditions, facing growing pressures from climate change, diseases, invasive species, and pesticides. Well-designed breeding programs can support sustainable beekeeping and essential pollination services.

## Introduction

In France, structured honeybee breeding programs remain marginal. The country lacks a long-standing honeybee breeding culture, and beekeepers have historically relied on foreign breeders to acquire genetically improved queens. Recently, however, diverse small-scale groups have initiated efforts to build breeding programs, although these initiatives are still in their early stages and often lack formal structure. This situation is not unique to France. Although some exceptions exist (Hoppe et al., 2020; Brascamp et al., 2016), in most countries, honeybee breeding plans are relatively recent and varied, from small-scale initiatives (Kistler et al., 2024; Maucourt et al., 2021) to large-scale ones (e.g., Mandl, 2024). Furthermore, beekeepers are particularly diversified in their motivations to keep bees, compared to other animal farmers. A large proportion are hobbyists who keep bees for varied non-profit related reasons (see, e.g., Chauzat et al., 2013, for Europe; Kulhanek et al., 2017, for the USA; Zalilova et al., 2021, for Russia). Even among professionals, considerable variation exists in practices, production environments, and the genetic backgrounds of their colonies.

Yet, little is known about what beekeepers themselves would consider relevant breeding goals. To date, only two studies were dedicated to this question, each focusing on a small group of hobbyists and a limited set of traits (Brascamp and Van Der Lans, 2025; Guichard et al., 2019). In addition, the European EurBeST study (Büchler et al., 2022) also provided information on breeding goals, by showing the expectation and level of satisfaction of queen buyers across Europe via an online questionnaire. However, these studies were limited to a few historical traits of common interest, and did not explore the possible variability in breeding goals across diverse beekeeping profiles within a country. Furthermore, Guichard et al.’s (2019) and Büchler et al.’s (2022) studies were part of approaches related to the selection of resistance to varroa mites, which influenced the questions in their questionnaires and their subsequent analyses.

In this context, we therefore explored the nature and diversity of breeding goals desired by French beekeepers according to their beekeeping profile. The beekeeping profile was characterized mainly by beekeeper’s level of professionalization, years of experience, main production, possible quality certifications, geographic location, and their bee’s genetic background. We inquired about which traits beekeepers wish to prioritize in their breeding goal based on a list of 20 possible traits, and what desired genetic change they would like for each trait of interest. To obtain a large and national panel of beekeepers, this study is based on results from an online survey directed to all French beekeepers we could reach, including beekeepers not directly involved in selective breeding.

Our study addresses two main research questions:

1. What traits do French beekeepers prioritize for breeding goals?
2. Would they generally agree with the same trait priorities regardless of their beekeeping profile?

## Material and methods

### Online Survey via a questionnaire

#### Creation and distribution of the survey questionnaire

An online survey was opened from mid-February 2023 to the last day of 2023. Its design had been informed by a preliminary study (Lauvie et al., 2024) in which a small number of beekeepers were surveyed with open questions that helped determine the relevant closed questions for the current study. The questionnaire was in French and implemented with Limesurvey (LimeSurvey Project Team and Schmitz, 2012). A text transcription of the questionnaire is available in the Supplementary Material S1, in French as well as in an English translation. The web link to the survey was distributed via the diverse media of several regional and national French beekeeping organizations, mainly: the INRAE-ITSAP mixed-technology unit ETTAP (previously PrADE until 2025), the French national queen breeders’ association (ANERCEA), the regional associations for the development of professional beekeeping (ADAs), and the French association of royal jelly producers (GPGR).

#### Survey structure and general content

The questionnaire was organized in three parts. The first inquired on the respondent’s beekeeping profile, the second on the characteristics of their bee stock, and the third on their ideal breeding goal. In the first part, respondents answered 10 general questions about their beekeeping profile (e.g., geographic area, years of experience, main products, quality labels), including also the genetic background of their bee stock. The following choices were proposed for the genetic background: ‘black (*mellifera*)’; ‘Carnolian (*carnica*)’; ‘Caucasian (*caucasica*)’; ‘Italian (*ligustica*)’; ‘local, undetermined, no particular control’; ‘Buckfast’; ‘royal jelly-specialized population’; ‘other’, accompanied by a free text field. This first part ended by asking for how many herds (one to three possible) the respondents wished to express breeding goals for. In the second part, respondents were asked about specifics of the herd they wished to express breeding goals for, informing about the herd’s size (i.e. number of colonies), whether these colonies were mostly transhumant or sedentary, and queen-rearing practices (if any). The third part addressed breeding goals. Respondents were first asked to rate the importance of including each trait in a breeding goal, and then to specify what direction of change was desired for each trait. To ensure accessibility to beekeepers beyond specialized breeders, we framed the question in practical terms: “Indicate a degree of usefulness for each of the traits listed below, according to how important it is for you to consider these traits in genetic choices for herd renewal.” Twenty potential traits were listed. Respondents rated each on a five-level scale: ’Useless’, ’Little useful’, ’Useful’, ’Very useful’, or ’Essential’, while ’Without response’ was the default answer. A warning message reminded respondents that “the more useful traits you include in the breeding goal, the more difficult it will be to improve strongly each of them.” For traits not rated as ’Without response’ or ’Useless’, a follow-up question asked respondents to specify the desired genetic trend using a four-level scale: ’Increase’, ’Decrease’, ’Maintain’, or ’Eliminate severe defects’.

### Data curation

#### Filtering duplicates and elimination of responses of untargeted areas

Of the 263 initial questionnaire answers, nine were excluded because they were duplicates from the same beekeepers. Duplicates were identified by near-identical answers leaving identical contact details. In these cases, the most recent answer was kept. As the study focused on breeding goals, five additional responses were excluded because the respondents reported no production or service activity (including non-monetary ones) related to beekeeping. Last, six other answers were excluded because they came from outside metropolitan French regions and were in too limited numbers to be informative (three from various French overseas territories and three from Algeria). As a result, 243 answers were kept, including 30 from respondents who provided information for 2 different herds with distinct breeding goals, leading to 273 combinations of breeding goal traits priorities listed in total.

#### Categorization and standardization

To describe beekeepers’ profiles, we derived secondary categorical variables by harmonizing heterogeneous free-text responses and, where necessary, aggregating categories to ensure sufficient sample sizes for statistical analyses. French administrative regions and free-text geographic responses were grouped into broader areas with internationally recognizable names, based primarily on geographic proximity and ecological context.

Similarly, main and secondary production activities were grouped into composite categories reflecting increasing diversification of outputs (e.g. honey only; honey and other hive products; honey and queens or swarms; honey, pollination, and other hive products). Two atypical responses, of beekeepers producing no honey at all, did not perfectly match any category but were assigned to the closest relevant group: one respondent reporting only queen or swarm production was grouped with *honey and queens or swarms*, and one respondent reporting only pollination was grouped with *honey and pollination and other hive products*.

For the herd genetic background, free-text entries were matched to one of the proposed categories whenever possible. Furthermore, when respondents reported raising multiple genetic backgrounds but subsequently described working with a single population and provided a single breeding goal, likely reflecting a heterogeneous genetic background, we assigned these responses to a new category: ‘various breeds & hybrids’. Because of their low number, two free-text responses referring to the *carpatica* subspecies and five to the *caucasica* subspecies were also assigned to this category. A last category was created as ‘breeder populations’ to gather six responses listing a free-text answer referring to a breeder’s name and 19 responses referring to breeding groups. These latter respondents had initially selected the responses ‘other’, ‘local, undetermined, no particular control’, or had listed multiple breeds but described working with a single population and provided a single breeding goal.

We also reassigned 21 respondents to the category ‘not involved in selective breeding’ who reported conducting selective breeding individually (outside a breeding group) but did not perform (or commissioned) any form of queen rearing. Not having the possibility, in such an online survey, to explore further why respondents considered themselves as engaged in selective breeding, we chose to rest upon a usual norm in genetics. Indeed, it is a common view to consider that, without participation in a breeding group, and thus when having to perform all breeding steps individually, performing queen rearing is an essential practice to be regarded as actively engaged in selective breeding. Similarly, we reclassified as ‘not actively engaged in breeding’ three respondents who indicated performing their own selective breeding, but actually relied on purchased dam queens to rear offspring queens as their production queens, which is a multiplication activity rather than selective breeding one in usual breeding scheme terminology.

The survey proposed six levels of importance for each trait to be included in a breeding goal (as detailed in section *Survey structure and general content*). However, the interpretation of the intermediate levels appeared to be inconsistent as respondents did not use the full range of six levels in a homogeneous way. We therefore merged the categories ‘Without response’, ‘Useless’, and ‘Little useful’ into a single ‘Useless’ level, and the categories ‘Useful’ and ‘Very useful’ into a single ‘Useful’ level, resulting in three final levels of importance: ‘Useless’, ‘Useful’, and ‘Essential’.

For the same reason, the two categories of the desired direction of genetic change ‘Eliminate severe defects’ and ‘Maintain’ were merged into a unique ‘Maintain’ category. Furthermore, when respondents wished genetic values to be directionally changed, less than 2% wanted them to decrease. Minor exceptions concerned tendency to propolize (4% for decreasing), and the absence of wax bridges between frames (9% for decreasing), the latter however likely reflecting a misinterpretation caused by the double negation created when applying the response ‘diminishing’ to the negatively defined trait ‘absence of wax bridges’ (Supplementary Material S1). As opposite directions of change were very marginal, the categories ‘Increase’ and ‘Decrease’ were grouped into a single ‘Improve’ category. Consequently, the initial four categories were reduced to only two: ‘Maintain’ and ‘Improve’.

### Statistical Analysis of data

Data analyses were run in R v4.5.1 (R Core Team, 2025) using the packages ‘data.table’ (Barrett et al., 2025) and ‘tidyverse’ (Wickham et al., 2019) for general data formatting and figure production, and ‘rvg’ (Gohel, 2025) and ‘officer’ (Gohel and Moog, 2025) for figure exportation. Multiple correspondence analysis (MCA) was performed to explore associations among beekeeper profile variables. The analysis was conducted using the ‘FactoMineR’ package (Lê et al., 2008) in R.

The relationships between the importance scoring of traits in the breeding goal and respondent’s beekeeping profile variables were analyzed by ordinal multiple regression models using the cumulative link model (CLM) function implemented in the ‘ordinal’ package (Christensen, 2023) in R (R Core Team, 2025). We used the default logit link function, which corresponds to the Proportional Odds Model (or Ordered Logit Model). The CLM function is appropriate for ordinal response variables as it accounts for the ordered nature of importance ratings while avoiding assumptions of equal intervals between response categories.

Initial ordered logit models included all explanatory variables (genetic background, production category, level of professionalization, experience, colony transhumance, geographic area of operations, or involvement in collective breeding) in a full model for each response variable, i.e. for the importance rating of each trait to be included in the breeding goal. Starting from the full model for a given response variable, reduced models were derived by sequentially removing variables with p-values greater than 0.15, based on likelihood ratio tests using the drop1 function from the ‘stats’ package in R (R Core Team, 2025), followed by refitting the model. Remaining variables were further assessed for statistical significance, and models were compared using the Akaike Information Criterion (AIC). The final model was selected as the one with the lowest AIC (or close to the lowest value) and *P*-values strictly inferior to 0.05 for the remaining explanatory variables.

## Results

### Survey respondent’s beekeeping profiles

Respondents covered all experience levels, based on years of beekeeping activity, approximately homogeneously (Table 1). About 20% of respondents were hobbyists, while 61% belonged to small to large professional operations, the rest being semi-professionals (Table 1). Geographical coverage generally aligned with France’s main beekeeping regions (FranceAgriMer, 2022), with few respondents from the Center (3%), North (5%) and Brittany (6%), intermediate levels of respondents from North-East (10%), South-West (10%) and West (15%), and larger proportions of respondents from East and South-East regions (25% each, Table 1).

**Table 1.** Number and proportion of respondents per years of experience, level of professionalization and main geographic area of beekeeping activity.

| Respondents' profile | Nb of respondents | Percentage (%) |
| --- | --- | --- |
| Years of experience <sup>1</sup> |  |  |
| Junior (0 to 4 years) | 56 | 23.0 |
| Intermediate (5 to 9 years) | 64 | 26.3 |
| Proficient (10 to 14 years) | 44 | 18.1 |
| Master (15 to 24 years) | 36 | 14.8 |
| Senior (25 years and more) | 43 | 17.7 |
| Level of professionalization |  |  |
| Hobbyist (no revenue) | 49 | 20.2 |
| Small semi-professional (1 to 100 colonies) | 21 | 8.6 |
| Large semi-professional (101 to 280) | 24 | 9.9 |
| Small professional (1 to 200 colonies) | 36 | 14.8 |
| Medium professional (201 to 400 colonies) | 59 | 24.3 |
| Large professional (401 to 1800) | 54 | 22.2 |
| Geographic area <sup>2</sup> |  |  |
| North | 13 | 5.3 |
| North-East | 24 | 9.9 |
| East | 61 | 25.1 |
| Center | 8 | 3.3 |
| South-East | 60 | 24.7 |
| South-West | 26 | 10.7 |
| West | 37 | 15.2 |
| Brittany | 14 | 5.8 |

<sup>1</sup> Years of experience: for hobbyists, i.e. beekeepers declaring no income from their activity, the anteriority was based on the period since starting beekeeping; for semi-professionals and professionals, it was based on the period since founding their beekeeping company.
<sup>2</sup> Geographic area: North stands for answers indicating Hauts-de-France and Belgium; North-East for Bourgogne-Franche-Comté and Grand-Est; East for Auvergne-Rhône-Alpes and Switzerland; Center Centre-Val de Loire, Île-de-France, and Normandy; South-East for Provence-Alpes-Côte d'Azur; West for Nouvelle-Aquitaine and Pays de la Loire; Brittany for Brittany.

All respondents, except two, reported producing at least honey, with 23% indicating it as their sole production (Table 2). In addition to honey, 10% declared to produce royal jelly as a main or secondary production, 7% to engage in pollination services, and 46% to produce queens or swarms (Table 2). Regarding quality labels, 38% were certified at least as organic, while 11% held other labels such as ‘Protected Geographic Indication’, or the French specific labels ‘Label Rouge’ or ‘Nature et Progrès’ (Table 2). Regarding colony mobility in a herd for which a breeding goal was provided, 29% of respondents reported sedentary colonies, 29% reported mixed itineraries (part sedentary, part transhumant), and 42% fully transhumant herds (Table 2). Last, of all survey respondents, 81% declared being actively involved in selective breeding activities, with 19% of all respondents doing so in a collective breeding group (Table 2).

**Table 2.**
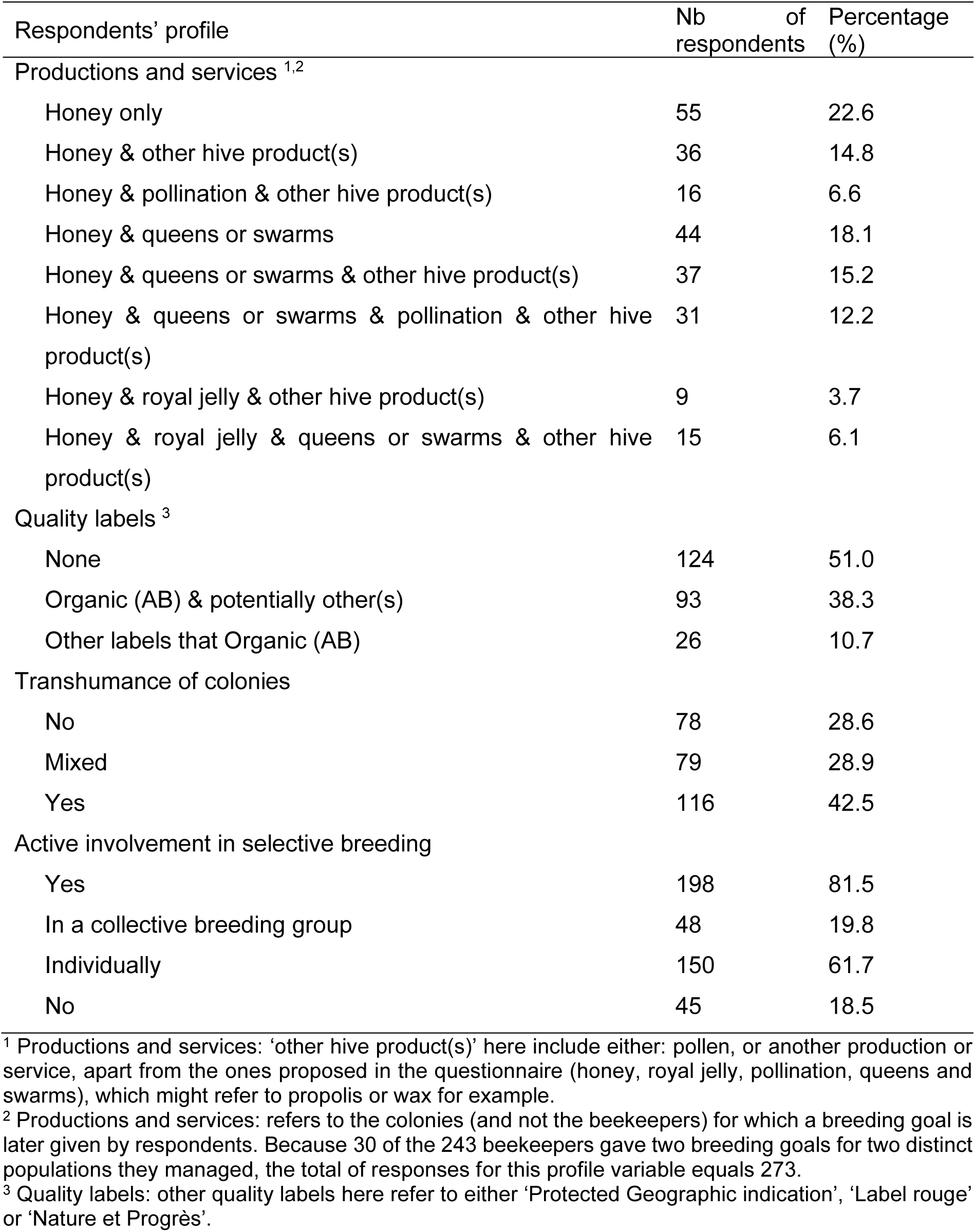
Number and proportion of respondents per type of productions, quality label, transhumance of colonies, and involvement in a breeding group.

| Respondents' profile | Nb of respondents | Percentage (%) |
| --- | --- | --- |
| Productions and services <sup>1,2</sup> |  |  |
| Honey only | 55 | 22.6 |
| Honey & other hive product(s) | 36 | 14.8 |
| Honey & pollination & other hive product(s) | 16 | 6.6 |
| Honey & queens or swarms | 44 | 18.1 |
| Honey & queens or swarms & other hive product(s) | 37 | 15.2 |
| Honey & queens or swarms & pollination & other hive product(s) | 31 | 12.2 |
| Honey & royal jelly & other hive product(s) | 9 | 3.7 |
| Honey & royal jelly & queens or swarms & other hive product(s) | 15 | 6.1 |
| Quality labels <sup>3</sup> |  |  |
| None | 124 | 51.0 |
| Organic (AB) & potentially other(s) | 93 | 38.3 |
| Other labels than Organic (AB) | 26 | 10.7 |
| Transhumance of colonies |  |  |
| No | 78 | 28.6 |
| Mixed | 79 | 28.9 |
| Yes | 116 | 42.5 |
| Active involvement in selective breeding |  |  |
| Yes | 198 | 81.5 |
| In a collective breeding group | 48 | 19.8 |
| Individually | 150 | 61.7 |
| No | 45 | 18.5 |

<sup>1</sup> Productions and services: 'other hive product(s)' here include either: pollen, or another production or service, apart from the ones proposed in the questionnaire (honey, royal jelly, pollination, queens and swarms), which might refer to propolis or wax for example.
<sup>2</sup> Productions and services: refers to the colonies (and not the beekeepers) for which a breeding goal is later given by respondents. Because 30 of the 243 beekeepers gave two breeding goals for two distinct populations they managed, the total of responses for this profile variable equals 273.
<sup>3</sup> Quality labels: other quality labels here refer to either 'Protected Geographic indication', 'Label rouge' or 'Nature et Progrès'.

Table 3 shows the number of answers attributed to each genetic background category. The least represented genetic background categories were *carnica* (6%), the royal jelly specialized population (7%), and *mellifera* and breeder populations (both at 9%), while the most represented ones were the Buckfast and the ‘various breeds & hybrids’ categories (each representing 26% of answers), the remainder 16% responses corresponding to the category local.

**Table 3.** Number and proportion of responses per bee genetic background category.

|  | Nb of responses | Percentage (%) |
| --- | --- | --- |
| Number of bee genetic backgrounds declared per respondent <sup>1</sup> |  |  |
| 1 | 142 | 58.4 |
| 2 | 76 | 31.3 |
| 3 | 22 | 9.1 |
| 4 | 3 | 1.2 |
| Total number of responses per bee genetic background |  |  |
| *carnica* | 16 | 5.9 |
| *mellifera* | 24 | 8.8 |
| Buckfast | 72 | 26.4 |
| royal jelly specialized population | 19 | 7.0 |
| local | 45 | 12.8 |
| breeder populations <sup>2</sup> | 25 | 9.2 |
| various possible breeds & hybrids <sup>3</sup> | 72 | 26.4 |

<sup>1</sup> Number of breeds declared per respondent: as declared by respondents before possible reattribution to categories ‘breeder populations’ or ‘various possible breeds & hybrids’.
<sup>2</sup> breeder populations: breeder or breeding group’s populations, excluding the national breeding group for the royal jelly producing population.
<sup>3</sup> various possible breeds & hybrids: responses listing multiple breeds but subsequently describing working with a single population and providing a single breeding goal.

### Multiple component analysis of survey respondent’s beekeeping profiles

Fig. 1 shows the multiple correspondence analysis (MCA) of respondent beekeeping profile variables, depicting how beekeeping characteristics clustered together.

**Fig. 1.**
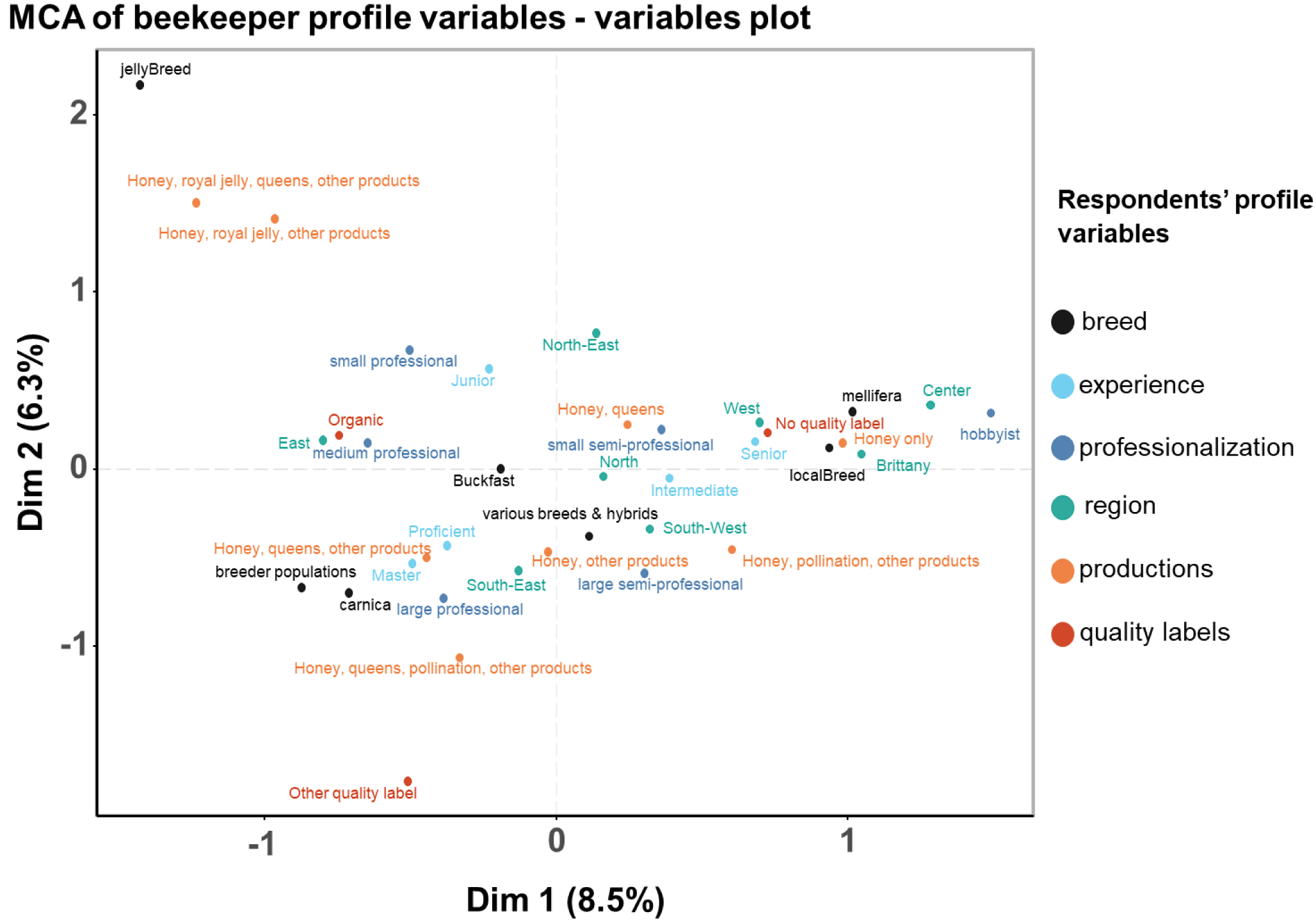
Graph of variables of the multiple correspondence analysis of beekeeper’s profile variables. Percentages on the axis titles indicate the proportion of total inertia explained by each axis.

The variables colony transhumance and collective breeding involvement were excluded to improve readability and because variables with fewer levels can contribute less than proportionally to inertia in MCA when analyzed alongside variables with more levels. The analysis reveals several distinct profile clusters. The cluster composed by the majority of respondents (see the middle of Fig. 1) comprised semi-professionals from the North, South-West, or South-East, that produce honey and queens or swarms, or honey and other hive products using Buckfast or various breeds & hybrids.

Royal jelly producers with royal jelly specialized populations formed a unique cluster distant from the origin, reflecting both their distinct profile and their low proportion in the overall set of respondents. Hobbyists from Center, Brittany and West regions, producing only honey with local or *mellifera* bees, and without quality labels, clustered on the right side of the first axis. Organic, medium-professional beekeepers from the East region formed an opposing cluster on the left. A last cluster comprised experienced beekeepers (10-24 years), large professionals with diversified productions (honey, queens or swarms, other products, and possibly also engaging in pollination services), and that keep *carnica* or breeder populations’ bees.

### Global ranking of traits in breeding goals

Across all responses, the median number of traits considered as at least useful was 13, while the median number of traits given as essential was 3, both indicating that ideal breeding goals are multi-trait for a majority of respondents (see Supplementary Fig. S1 for the distribution of the number of useful (a) or essential (b) traits per respondent). About a quarter of all respondents reported that the combination of the 3 traits: honey yield, disease resistance, and swarming tendency, was essential to include in the breeding goal (Supplementary Table S1). Fig. 2 shows the proportion of beekeepers that considered a trait as useful or essential to include in the breeding goal (a), and the proportion that wished a trait to be either improved or maintained at its current level of performance (b).

**Fig. 2.**
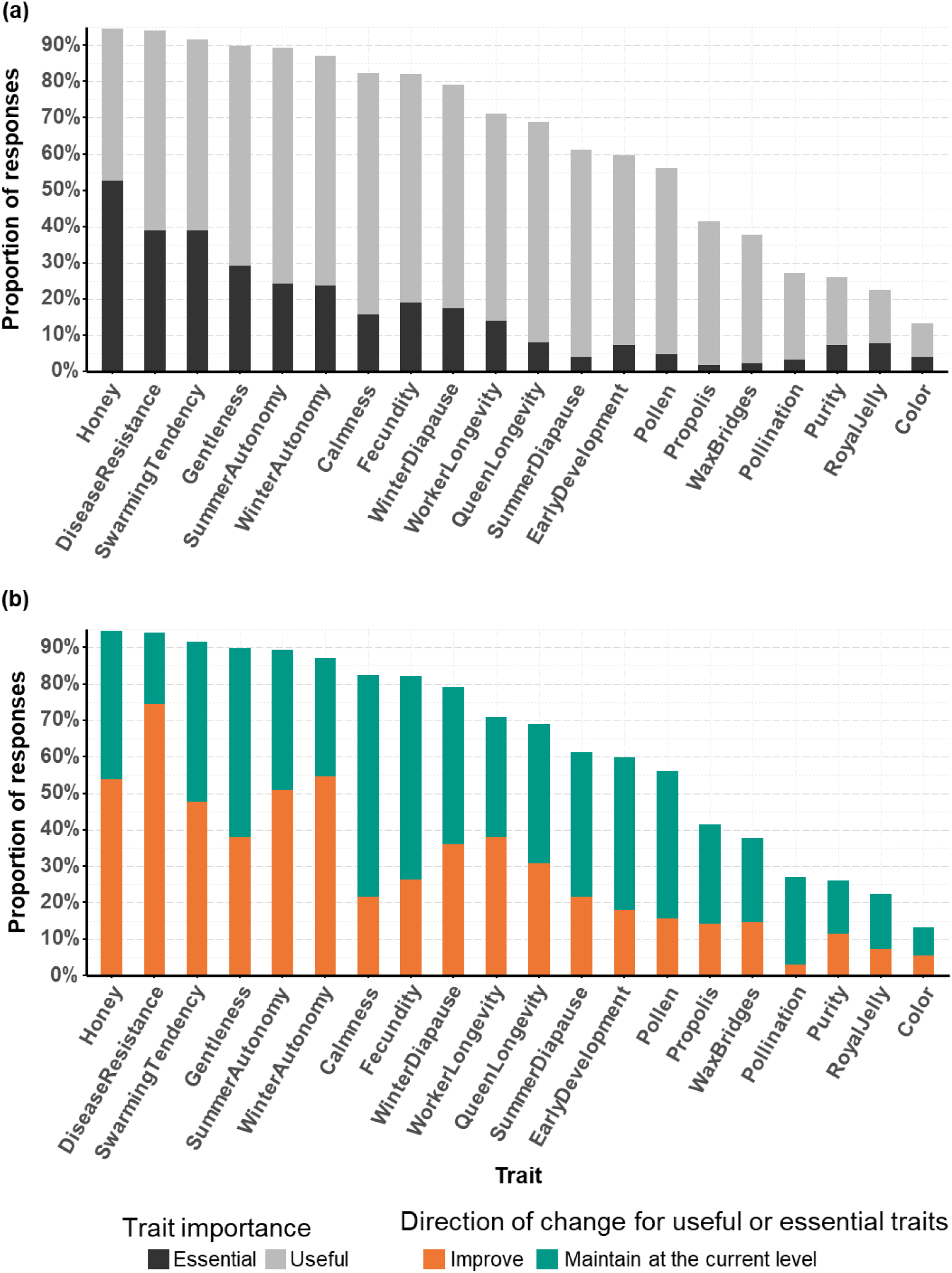
Proportion of responses indicating each trait as useful or essential to include in the breeding goal and desired direction of change. Honey: honey yield; DiseaseResistance: disease resistance (in the broad sense); SwarmingTendency: swarming tendency; Gentlness: propensity of bees to not attack beekeepers during colony visit; SummerAutonomy and WinterAutonomy: resp. summer and winter feed autonomy (no need of feeding); Calmness: propensity of bees to remain calm on the frames during colony visit, and not to run and cluster together; WinterDiapause and SummerDiapause: resp. winter and summer diapause (reduced colony activity, including egg-lay reduction or absence); fecundity: brood quantity produced in the colony; WorkerLongevity and QueenLongevity: resp. worker and queen longevity; pollen: pollen production; propolis: propolis production; WaxBridges: propensity to build wax bridges between frames; Pollination: pollinating capacity; BreedPurity: breed purity; JellyYield: royal jelly yield.

Almost all respondents (approximately 94%, Fig. 2a) considered honey yield and disease resistance to be at least useful to include in the breeding goal, with honey yield being more frequently rated as essential (53% vs 39%). However, disease resistance was the trait most beekeepers wished to improve (74%, Fig. 2b) rather than to only maintain it at its current level, while only 54% of beekeepers wished to improve honey yield. The other traits considered at least useful and to be improved for a majority of respondents included winter (55%) and summer (51%, Fig. 2b) feed autonomy.

Over 85% of beekeepers considered swarming tendency and feed autonomy (both in summer and winter) as at least useful, and about 80% or more also indicated the traits gentleness, calmness, fecundity, and winter diapause, as at least useful (Fig. 2a). While about half of the respondents wished to improve swarming tendency and summer feed autonomy, less than 40% wished to genetically improve gentleness, calmness, fecundity, and winter diapause (Fig. 2b). Traits with less than 20% of respondents wishing to improve them included earliness of colony development, pollen production, propolis production, and pollination (ranking lowest, at 3%, Fig. 2b).

### Influence of beekeeping profiles on breeding goal choices

Table 4 presents the significance of the main respondents’ beekeeping profile variables on the perceived importance of including each trait in the breeding goal, based on the best-fitted ordinal multiple regression model for each trait. Results focused on the 9 traits considered as useful or essential to include in the breeding goal by more than 75% of respondents (Fig. 2). The corresponding table for remaining traits, of secondary interest, is available as Supplementary Table S2.

**Table 4.** Beekeeping profile variables significantly influencing the importance attributed to the main traits of interest in the breeding goal, based on fitted ordinal multiple regression models.

| Trait | AIC |  | Profile variables |  |  |  |  |  |  |
| --- | --- | --- | --- | --- | --- | --- | --- | --- | --- |
|  | Full model | Best fitted model<sup>1</sup> | Bee genetic background | Organic certification | Productions | Experience | Level of professionalization | Colony sedentariness | Area |
| Honey yield | 434.87 | 419.77 | *** | * |  |  | * | * |  |
| Disease resistance | 499.50 | 467.58 |  |  | ** |  |  |  |  |
| Swarming tendency | 481.26 | 467.79 | *** |  |  | * |  |  |  |
| Gentleness | 464.08 | 445.51 | *** |  |  |  |  |  |  |
| Calmness | 466.40 | 431.37 | *** | *** |  |  |  |  |  |
| Summer autonomy | 498.28 | 470.03 |  |  |  |  |  |  | * |
| Winter autonomy | 504.95 | 475.31 | ** |  |  |  | * |  |  |
| Fecundity | 522.03 | 497.31 |  | ** |  |  |  |  |  |
| Winter diapause | 526.47 | 507.49 | ** |  |  |  |  |  |  |

Level of significance: (\*): $0.01 < p.\text{value} < 0.05$ ; (\*\*): $0.001 < p.\text{value} < 0.01$ ; (\*\*\*): $p.\text{value} < 0.001$ .
AIC Full model: Akaike information criterion of the ordinal multiple regression model including all profile variables to explain the importance attributed to a trait in the breeding goal.
AIC Best fitted model: Akaike information criterion of the model including only statistically significant profile variables as explanatory variables.
<sup>1</sup> The best fitted ordinal multiple regression model is the model with the lowest AIC compared to the full model after model reduction using likelihood ratio tests to remove non-significant profile variables.

#### Effects of bee’s genetic background on the degree of importance of trait inclusion in the breeding goal

Bees’ genetic background significantly influenced the importance attributed to six out of the nine traits of main interest (Table 4). Fig. 3 shows the distribution of importance ratings across genetic backgrounds for the six traits for which the genetic background had a significant effect on traits’ importance rating.

**Fig. 3.**
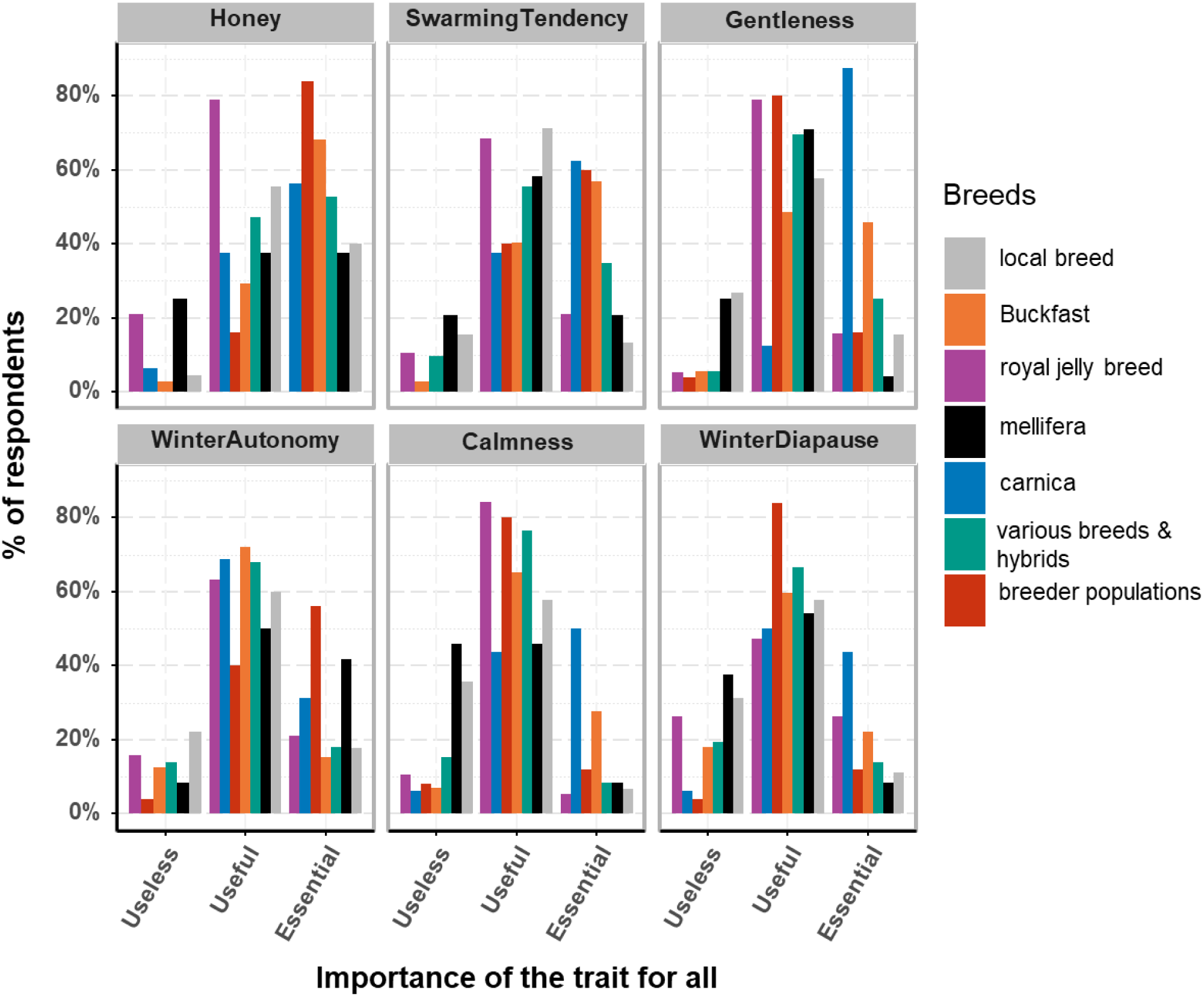
Proportion of responses indicating each trait as useless, useful, or essential to include in the breeding goal per bees’ genetic background. Honey: honey yield; SwarmingTendency: swarming tendency; Gentlness: propensity of bees to not attack beekeepers during colony visit; WinterAutonomy: winter feed autonomy (no need of feeding); Calmness: propensity of bees to remain calm on the frames during colony visit, and not to run and cluster together; WinterDiapause: winter diapause (reduced colony activity, including egg-lay reduction or absence).

In Supplementary Table S3, exact proportions of these importance rating frequencies for all traits are also reported. Statistically significant contrasts in the fest-fitted CLM (Table 4) will be illustrated by mentioning the differences in raw scoring frequencies. For honey yield, keepers of royal jelly-specialized bees rated ‘essential’ this trait in a lower proportion (0%) than keepers of breeder populations (84%), Buckfast (68%), and various breeds & hybrids (53%). *Mellifera* keepers also rated ‘essential’ honey yield in a lower proportion (38%) than keepers of breeder populations and Buckfast. For swarming tendency, keepers of *carnica* (63%), breeder populations (60%), and Buckfast (57%), rated ‘essential’ this trait at similarly high proportions, while keepers of royal jelly-specialized (21%), *mellifera* (21%), and local bees (13%) rated it significantly lower. Gentleness showed a pronounced breed-dependent variation (Fig. 3). *Carnica* keepers rated it essential the most often (88%), attributing it statistically more importance than all others except Buckfast and royal jelly-specialized keepers: only 16% of local breed or of breeder populations keepers rated it essential, and 4% of *mellifera* keepers. Still regarding gentleness, Buckfast keepers rated it essential more often (46%) than *mellifera* and local breed keepers. Various breeds & hybrids keepers rated it essential less often than *mellifera* keepers. For winter feed autonomy, breeder population keepers rated it essential (56%) more often than Buckfast (15%) and various breeds & hybrids (18%) keepers. Last, for winter diapause, *carnica* keepers rated this trait more often as essential (44%), than mellifera (8%) and local breed (11%) keepers.

#### Effects of organic certification, transhumance practice, farm region and productions on breeding goal preferences

The other beekeeping profile variables: organic certification, farm productions, transhumance practice, and geographic area, each significantly influenced the importance of one to three traits out of the nine traits of main interest (Table 4), while the variable ‘active involvement in selective breeding’ never significantly explained the importance attributed to any of these traits.

Organic certification significantly influenced importance ratings for honey yield, fecundity, and calmness (Table 4, Fig. 4). In all three cases, certified organic beekeepers rated these traits as less essential (50%, 11%, and 5%, respectively, Fig. 4 and Supplementary Table S4) compared to non-certified beekeepers (55%, 25%, and 23%, respectively). While transhumance practice also significantly impacted the importance of honey yield, specific pairwise contrasts were never significant. However, beekeepers practicing at least partial transhumance tended to attribute more importance to honey yield than those with sedentary colonies. The importance attributed to summer feed autonomy was only influenced by the region with Brittany-based beekeepers tending to rate it as more essential than the others. Noteworthy, the area did not significantly affect the importance rating of other traits that could be expected as variably problematic across the diverse French biotopes, such as winter feed autonomy or winter diapause. Last, farm productions significantly influenced only disease resistance importance, with beekeepers producing honey, queens or swarms and other hive products, rating it more essential (60%) than beekeepers producing honey with or without queens or swarms (30%), or with royal jelly and other products (13%, Supplementary Table S4).

**Fig. 4.**
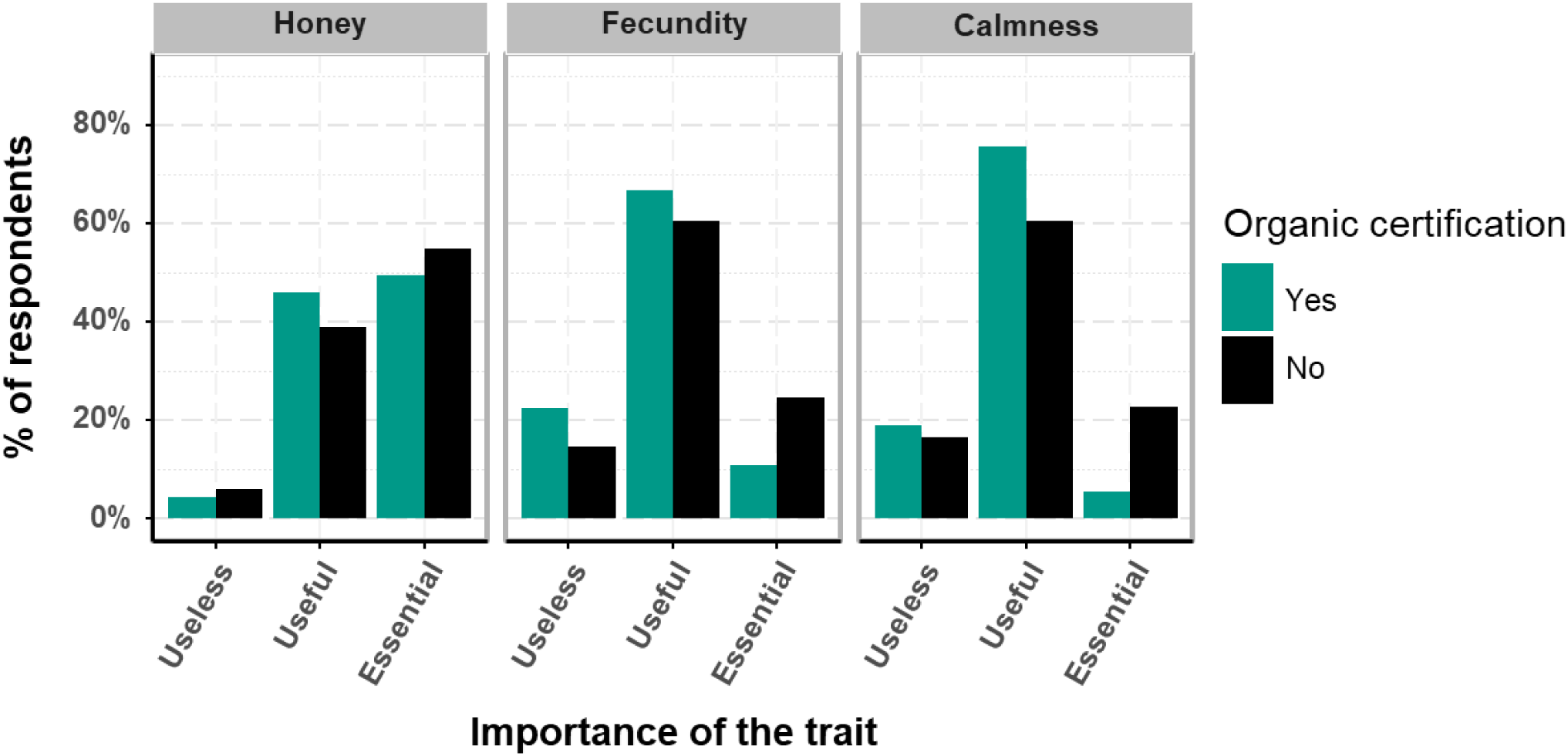
Proportion of responses indicating each trait as useless, useful, or essential to include in the breeding goal per organic certification status. Honey: honey yield; fecundity: brood quantity produced in the colony; Calmness: propensity of bees to remain calm on the frames during colony visit, and not to run and cluster together.

#### Effects of experience and professionalization level of beekeepers on breeding goal preferences

The beekeeping profile variables ‘experience level’ and ‘professionalization level’ significantly influenced the importance rating of only one or two traits out of the nine traits of main interest (Table 4). Experience significantly influenced only the importance given to swarming tendency, with the least experienced beekeepers (juniors, < 5 years) rating it more frequently as essential (51%) than proficient ones (10-14 years, 24%, and Supplementary Table S4). As the professionalization level increased, the importance attributed to honey yield tended to increase, with large semi-professionals (56%), and small (50%) and large (69%) professionals rating it more essential than hobbyists (28%, Supplementary Table S4). Small (32% essential) and medium (31%) professionals also attributed more importance to winter autonomy compared to hobbyists (10%, Supplementary Table S4).

## Discussion

This study, based on about 250 answers from mainly professional beekeepers to an online questionnaire, documents the numerous traits useful to integrate in a breeding goal according to French beekeepers and how they vary according to beekeeping profiles. First, it highlights that beyond the usual traits targeted in bee breeding plans, such as honey production, disease resistance, swarming tendency and gentleness, other traits of frequent interest, such as feed autonomy in summer and in winter, are poorly studied. Second, it reveals that the main driver that influences trait priorities in the breeding goal is the bees’ genetic background. Third, it shows that beekeepers generally consider that relevant breeding goals would be clearly multitrait, raising questions about the feasibility, relevance, and practical implementation in breeding programs, including questions on how to reach consensus on breeding goal definition in the case of participatory approaches in breeding groups.

### A shared global breeding goal across all beekeeping profiles

Respondents appeared to globally envision similar breeding goals. To practically all respondents, honey yield and disease resistance appeared as universally important traits. A vast majority wished to improve disease resistance while a lesser majority wished to improve honey yields. This widens previous results studying specific hobbyists breeding groups (Brascamp and Van Der Lans, 2025; Guichard et al., 2019), and complements results reported in the European EurBeST project (Büchler et al., 2022), all showing disease resistance as the trait to be improved for the most respondents. This major focus on improving disease resistance traits is also visible in the progressively growing addition (starting around the mid-2000s) of records in the largest honeybee breeding database (BeeBreed, Bienefeld and Hoppe, 2024) on the general brood health-related trait hygienic behavior (HYG) and the trait *Varroa* infestation development (VID). The perceived importance of health-related traits is also reflected by the increasing weight attributed to the varroa index in the published estimated breeding value for total merit in BeeBreed. Emerging breeding initiatives often also integrate these or similar disease resistance traits (Arista Bee Research, 2024; Kistler et al., 2024; Maucourt et al., 2023).

Apart from the bees’ genetic background, beekeeping profile variables had a very limited influence on breeding goals. In particular, while organic certified beekeepers attributed a significantly lower importance to honey yield, fecundity, and calmness, differences remained of a modest size (Fig. 4), so that organic certified beekeepers appeared to have compatible breeding goals with non-certified beekeepers, as found in previous individual surveys (Lauvie et al., 2024) as well as in two collective workshops with certified and non-certified beekeepers together (Lauvie et al., 2026).

### A major impact of bees’ genetic background on breeding goal trait priorities

By far, the profile variable with the largest influence on breeding goal trait prioritization was the bees’ genetic background. Although France’s native honeybee is the *mellifera* subspecies (Ruttner, 1988), a large diversity is present nowadays (Wragg et al., 2022), allowing French beekeepers to choose breeds that best suit their production objectives. Our results revealed a general alignment between the reputed characteristics of certain genetic backgrounds and the traits their keepers prioritized. For example, as described by Ruttner (1988) and popularized to French beekeepers by Peyvel (1994), *mellifera* bees have a reputation of not being gentle nor calm, with a variable swarming tendency, some ecotypes swarming abundantly. In our study, *mellifera* keepers rated these three behavioral traits with an unfavorable reputation for *mellifera* as relatively unimportant compared to keepers of other genetic backgrounds (Fig. 3, Supplementary Table S3). Conversely, *carnica* bees are typically described as overwintering very well, in part due to a relatively longer winter diapause, and being among the gentlest and calmest bees (Peyvel, 1994; Ruttner, 1988). Accordingly, *carnica* keepers rated winter diapause, gentleness, and calmness, as the most important traits compared to other genetic backgrounds (Fig. 3, Supplementary Table S3).

This pattern underlines the diverse links between the traits to be integrated in the breeding goal for future evolution of the population, and the traits judged by beekeepers as important in their production system and promoted in the present by using a genetic background with a favorable reputation for these traits. For example, *carnica* keepers considered swarming tendency as a more important breeding goal trait than other beekeepers while *carnica* has an unfavorable reputation for this trait (Peyvel, 1994; Ruttner, 1988). Furthermore, the proportion of *carnica* keepers that would like to improve these four traits of interest compared to maintain them at their current level was among the lowest for the well-reputed traits (winter diapause, gentleness, calmness), but among the highest for the bad-reputed trait (swarming tendency) (Supplementary Fig. S2). This suggests that beekeepers did distinguish between maintaining desirable breed characteristics and voluntarily improving problematic ones.

### Future research priorities: from trait preferences to practical operational selection criteria

#### Translating breeding goal trait preferences into breeding goal trait weights

Our findings highlight breeding goal priorities across the French beekeeping community, but translating these general preferences into concrete breeding programs requires substantial work. A breeding plan is designed for a given bee population. There will therefore be different breeding objectives depending on genetic backgrounds, even though production, disease resistance, and swarming tendency should remain shared objectives. Pragmatic breeding goals will need to consider the availability of relevant selection criteria that can be recorded at least on a substantial part of the breeding population. In addition, the trait prioritization will need to be specifically quantified for the bee population of interest to derive actual breeding goal trait weights.

To support the definition of the weights for the traits of interest to be included in the breeding goal and associated merit index, various approaches have been developed. Their will is to integrate economic, environmental, and social priorities, resting upon different principles (Nielsen et al., 2014; Sölkner et al., 2008; Olesen et al., 2000). A classical approach is the desired gains approach, where breeders define acceptable trade-offs between genetic gains across traits. This option could be implemented in a merit index using the results we got from our questionnaire by stating that maximum and similar genetic gains are searched for traits essential to improve, and 50% (for instance) of this maximum gain is searched for traits that are only useful to improve, while keeping constant the performances of all traits to be maintained. However, regarding decision-making process in a breeding group, moving from individual trait preferences to a collective breeding goal relevant to all participants, raises a broader challenge from a participatory approach perspective. While survey-based approaches provide valuable insight into the diversity of individual viewpoints, they cannot be directly translated into collective breeding decisions.

#### Finding operational selection criteria for traits of interest to a large part of beekeepers

Furthermore, several traits rated here as at least useful by a large majority (≥70%) of beekeepers remain poorly studied regarding their phenotypic measurability and genetic parameters. Feed autonomy (in summer and in winter), queen fecundity, winter diapause, and worker and queen longevity all warrant further research to propose reliable, practical selection criteria with good prospects to respond to selection. While genetic parameter estimates have recently been published for some of these traits, such as queen fecundity and summer feed autonomy (Kistler et al., 2024), larger datasets and more genetic backgrounds need to investigated. Winter diapause has the reputation to vary considerably phenotypically within the species (Ruttner, 1988), and might be assessed remotely using brood temperature sensors, as in winter colonies cannot generally be visited (Godeau et al., 2023; Meikle et al., 2016). Such measurements might also be directly useful to beekeepers for management, indicating when acaricide oxalic acid treatments would be most effective, and identifying early on colonies without a brood-less period at the moment of treatment and thus potentially at risk of high varroa infestation in the following season. Worker longevity could potentially be estimated from brood surface and adult worker population data, needing investigation.

Disease resistance, including regarding varroosis, emerged as a high-priority trait, yet efficient selection criteria remain lacking despite numerous research efforts (Guichard, 2021). Identifying cheap, easily measured, criteria with high genetic coefficients of variation that reliably predict colony-level disease resistance remains a critical research need. Furthermore, as new threats emerge, such as *Paenibacillus melissococcoides* recently detected in diseased colonies in Switzerland (Ory et al., 2023, 2025), or that gain importance such as *Nosema ceranae* in Germany (Gisder et al., 2017), developing selection criteria for resistance to other pathogens also needs consideration. Advancing the phenotypic and genetic evaluation of these priority traits represents essential groundwork for sustaining developing breeding programs aligned with beekeeper needs.

## Conclusion

We gathered responses of about 250 French beekeepers, mainly semi and full-professionals, that shared their opinions on which traits matter most for honeybee breeding goals. Honey yield, disease resistance, swarming drive, and gentleness, emerged as the top-priority traits, followed by summer and winter feed autonomy, which ranked higher than the classically targeted trait calmness. Respondents envisioned clearly multitrait breeding goals. In operational breeding programs, fewer traits will probably be included compared to the stated ideals due to cost limitations and lack of suitable selection criteria. While breed purity appeared of little interest to French beekeepers, the genetic background the respondent used was the major source of variation in the importance attributed to each trait in their breeding goals, which suggests programs using bees of different genetic backgrounds would ideally use different breeding goals. Other beekeeper’s profile characteristics had only a marginal effect on breeding goal trait priorities. A noticeable part of respondents that considered a trait as useful to include in the breeding goal only wished the trait to be maintained at its current performance level, rather than to improve it. A major exception to this was disease resistance, to be improved genetically by about 75% of beekeepers. Several traits that appeared as at least useful to consider in a breeding goal by a large majority (∼70% or more) of beekeepers are not or only poorly studied regarding their measurability and potential to respond to selection, such as feed autonomy traits, queen fecundity, diapause in winter, and worker and queen longevity. Future research is thus needed to explore possible selection criteria for these traits and assess their potential for genetic improvement.

## Supporting information

Survey transcription, Supplementary tables and a supplementary figure

## Ethics approval

Not required.

## Data and model availability statement

The cured data (see section Data curation for details) are available at: https://doi.org/10.57745/F3CEDI.

## Declaration of generative AI and AI-assisted technologies in the writing process

The main author sporadically used Claude AI and OpenAI’s online chatbots to help rephrasing certain problematic sentences. DeepL was sometimes used for French to English translations and then often reformulated for field-specific key words.

## Acknowledgements

The authors are thankful to the many beekeepers that took the time and effort to respond to the online questionnaire, thus contributing the research data used in this study. We are also thankful to Coline Kouchner for initial inputs on the questionnaire’s content and help in disseminating the survey to beekeepers.

## Author contributions

TK, BB, and FP designed the questionnaire. TK implemented and managed the online survey, coordinated its dissemination to beekeepers, performed the formal analyses, curated the data, and produced the figures. BB disseminated the survey to beekeeping organizations. TK and BB contributed to the conceptualization, investigation, data curation, formal analysis, and writing of the original draft. FP supervised the work and contributed to methodological choices. AL and FP acquired funding, administered the project, and contributed to the work’s conceptualization. All authors contributed to the interpretation of the results, reviewed and edited the initial draft, and approved the final version.

