## Supplementary material for "Beekeepers’ preferences for honeybee breeding goals: a French case study": Survey transcription, Supplementary tables and a supplementary figure

### Supplementary File – for Online Publication Only

### Supplementary Material S1

The original text transcription of the questionnaire, being in French, cannot be hosted on the chosen preprint platform. To obtain it, please directly contact the corresponding author.

[English translation]

#### Start-up message

Dear beekeeper,

As part of an INRAE research project, in partnership with ITSAP-Institut de l'abeille, ADAPI, ADA Occitanie, ADANA and Agri Bio Ardèche, we are asking you to identify the various beekeeping traits that you feel are important to improve genetically in your herd, so that we can quantify your selection requirements.

This questionnaire takes about 5 to 10 min to complete.

#### Section 1: Your beekeeping profile

**Beekeeping is an activity you engage in as a:** **(obligatory)**
full-time professional / multi-active professional / I'm a hobbyist

**In which region do you do most of your beekeeping?** **(obligatory)**
Auvergne-Rhône-Alpes / Bourgogne-Franche-Comté / Bretagne/Centre-Val de Loire / Grand Est / Hauts-de-France / Ile-de-France / Normandie / Nouvelle-Aquitaine / Occitanie / Pays de la Loire / Provence-Alpes-Côte d'Azur / Other (please specify).

**How long have you been beekeeping (if you are a professional, when did you found your company)?** **(optional)**
Select any year

**Farm productions** **('Without answer' selected by default)**
Table of possible productions (honey / royal jelly / pollination / sale of queens and swarms / others) by importance for the farm (main activity / secondary / not at all / / Without answer) - several choices possible by importance (e.g. possible to have several main productions)

**Is any of your production certified:** **(optional)**
No/In conversion / Organic Agriculture (AB) / Nature et Progrès / Label Rouge / Designation of Origin / Other (specify).

**Number of colonies on the farm (number at wintering, full-size hives + half-sized hives):** **(optional)**
Field to enter a number

**Are you involved in a breeding program?** **(obligatory)**
Yes, in a group / Yes, internally / No / Other: (specify)

If 'Yes, in a group': **What is the name of this selection group? (e.g. GPGR, ADAPI testing network, CETA Api d'Oc...)** **(optional)**

**Do you raise a particular breed or ecotype on your farm (multiple choice possible)?** **(optional)**
/ local, undetermined, no particular control
/ Buckfast
/ Royal Jelly bee
/ Black (mellifera)
/ Carnolian (carnica)
/ Italian (ligustica)

/ Caucasian (caucasica)
/ Other (specify)

**Do you keep several types of bees for different purposes and uses (e.g. royal jelly and honey), and for which you would like to answer the second part of this questionnaire (on the desired breeding goals) separately? (obligatory)**

no / 2 different herds (or more)

→ conditional question for the rest: if “no”, simplified version with a single herd, if more: version with repetition of the following 2 parts (once for each of the 2 herds)

**Section 2: Description of herd and breeding goals [specific to a single herd].**

**For the first herd: description and breeding goals [specific to several herds].**

**For the other herd: description and breeding goals [specific to several herds].**

**Size of the part of the herd for which you wish to express a breeding goal (in number of queens) (optional)** [specific to a single herd].

Free field

**Which first herd are we talking about (optional)** [specific to several herds].

Free field

**Size of herd in question (in number of queens) (optional)** [specific to several herds].

Indicate size

**Your apiaries of this herd are: (obligatory)**

All (or almost all) moving / All (or almost all) sedentary / A mixture: some sedentary, others moving

**Do you breed queens (or have someone do it for you)? (obligatory)**

Yes / No

If ‘Yes’: **I graft (or have grafts made) mainly from strains: (obligatory)**

Mainly bought or exchanged / Mainly from my own stock / Both at the same time

If ‘No’: **I don't graft (or don't have grafting done) and: (obligatory)**

I buy most of my production queens / I let most of my colonies ripen themselves / A mixture of both strategies

### Section 3: Which traits do you think will be useful in making genetic choices for the renewal of your herd? [specific to a single herd]

### Which traits do you find useful in the genetic choices for the renewal of this first/second herd? [specific to several herds; Repeats 2 times in this case]

Indicate a degree of usefulness for each of the **traits** listed below, according to how important it is for you to **consider** these traits **in genetic choices for herd renewal.** You will **then** be asked whether you would like to **improve, reduce or maintain these traits at their current level.**

Help: the more useful traits you include in the breeding goal, the more difficult it will be to improve strongly each of them.

**Table with 20 characters and 5 levels of usefulness (‘not useful’, ‘not very useful’, ‘useful’, ‘very useful’, ‘indispensable’) (‘Without answer’ selected by default)**

/ Honey yield
/ Pollen production
/ Royal jelly production
/ Ability to pollinate
/ Gentleness
/ Frame stability
/ Low propensity to swarm
/ Tolerance and/or resistance to varroa mites and diseases
/ Early season development
/ Queen fertility / Brood quantity
/ Winter lay stop
/ Summer egg-laying cessation (in the absence of resources)
/ Winter food self-sufficiency (no need to feed in winter)
/ Self-sufficient feeding in season (no need to feed in season)
/ Queen longevity
/ Longevity of workers
/ Tendency to propolize
/ Absence of wax bridges (parasitic constructions between frames)
/ Color
/ Racial purity

All traits are then listed in a table with a utility level from “Little useful” to “Indispensable”.

**For useful traits to include in your breeding goal, you want to: (obligatory)**
A: increase performance level,
B: lower performance level,
C: maintain current performance level,
D: eliminate severe defects.

### End of questionnaire

**Other comments, information or feedback on this topic): (optional)**
Free field

**Your contact details (optional)**
If you wish, you can give us an e-mail address so that we can send you the results of

the survey, or come back to you to clarify certain answers. Your address will be kept strictly confidential.

Free field

**Supplementary Table S1** Frequency of trios of trait reported as essential to include in the breeding goals

| Trio of essential traits | Frequency |
|---|---|
| Honey yield, Disease resistance, Swarming tendency | 23.8% |
| Honey yield, Disease resistance, Gentleness | 16.8% |
| Honey yield, Gentleness, Swarming tendency | 16.8% |
| Disease resistance, Swarming tendency, Winter diapause | 16.1% |
| Disease resistance, Gentleness, Swarming tendency | 15.8% |
| Disease resistance, Summer feed autonomy, Swarming tendency | 15.8% |
| Disease resistance, Swarming tendency, WorkerLongevity | 15.8% |
| Disease resistance, Fecundity, Swarming tendency | 15.4% |
| Disease resistance, Swarming tendency, Winter feed autonomy | 15.4% |
| Disease resistance, Summer autonomy, Winter feed autonomy | 14.7% |

**Supplementary Table S2** Beekeeping profile variables significantly influencing the importance attributed to the traits of secondary interest in the breeding goal, based on fitting ordinal multiple regression models.

| | AIC | | Profile variables | | | | | | | |
|---|---|---|---|---|---|---|---|---|---|---|
| Trait | Full model | Best fitted model [1] | Bee genetic background | Organic certification | Productions | Experience | Level of professionalization | Colony sedentariness | Area | Collective breeding |
| Worker longevity | 550.95 | NA | | | | | | | | |
| Queen longevity | 511.23 | NA | | | | | | | | |
| Summer diapause | 459.45 | 445.6 | | * | * | | ** | | | |
| Early development | 497.16 | 472.89 | *** | * | | | | ** | | |
| Pollen | 488.32 | 453.93 | | | * | | | | | * |
| Propolis | 413.56 | 401.18 | * | ** | ** | | | | | |
| Wax bridges | 439.82 | NA | | | | | | | | |
| Pollination | 371.26 | 358.52 | ** | * | * | | | | ** | * |
| Purity | 387.96 | 360.87 | *** | | | | | | | |
| Royal jelly | 289.3 | 268.19 | *** | | | | | | | |
| Color | 245.91 | 233.83 | *** | | * | | | | | |

[1] Level of significance: (*): 0.01 < p.value < 0.05; (**): 0.001 < p.value < 0.01; (***): p.value < 0.001.

AIC Full model: Akaike information criterion of the ordinal multiple regression model including all profile variables to explain the importance attributed to a trait in the breeding goal.

AIC Fitted model: Akaike information criterion of the model including only statistically significant profile variables as explanatory variables. AN here refers to the situation in which no model provided a better fit by step-wise selection of only explanatory variables with a statistically significant effect.

[1] The best fitted ordinal multiple regression model is the model with the lowest AIC compared to the full model after model reduction using likelihood ratio tests to remove non-significant profile variables.

**Supplementary Table S3** Importance rating frequency per genetic background for honey, swarming tendency, and gentleness

| Trait | Genetic background | Importance rating | | |
|---|---|---|---|---|
| | | **Useless** | **Useful** | **Essential** |
| Honey yield | jellyBreed | 21.1 | 78.9 | 0.0 |
| | carnica | 6.2 | 37.5 | 56.2 |
| | breeder populations | 0.0 | 16.0 | 84.0 |
| | Buckfast | 2.8 | 29.2 | 68.1 |
| | various breeds & hybrids | 0.0 | 47.2 | 52.8 |
| | mellifera | 25.0 | 37.5 | 37.5 |
| | localBreed | 4.4 | 55.6 | 40.0 |
| Swarming tendency | jellyBreed | 10.5 | 68.4 | 21.1 |
| | carnica | 0 | 37.5 | 62.5 |
| | breeder populations | 0 | 40.0 | 60.0 |
| | Buckfast | 2.8 | 40.3 | 56.9 |
| | various breeds & hybrids | 9.7 | 55.6 | 34.7 |
| | mellifera | 20.8 | 58.3 | 20.8 |
| | localBreed | 15.6 | 71.1 | 13.3 |
| Gentleness | jellyBreed | 5.3 | 78.9 | 15.8 |
| | carnica | 0.0 | 12.5 | 87.5 |
| | breeder populations | 4.0 | 80.0 | 16.0 |
| | Buckfast | 5.6 | 48.6 | 45.8 |
| | various breeds & hybrids | 5.6 | 69.4 | 25.0 |
| | mellifera | 25.0 | 70.8 | 4.2 |
| | localBreed | 26.7 | 57.8 | 15.6 |
| WinterAutonomy | jellyBreed | 15.8 | 63.2 | 21.1 |
| | carnica | 0.0 | 68.8 | 31.2 |
| | breeder populations | 4.0 | 40.0 | 56.0 |
| | Buckfast | 12.5 | 72.2 | 15.3 |
| | various breeds & hybrids | 13.9 | 68.1 | 18.1 |
| | mellifera | 8.3 | 50.0 | 41.7 |
| | localBreed | 22.2 | 60.0 | 17.8 |
| Calmness | jellyBreed | 10.5 | 84.2 | 5.3 |
| | carnica | 6.2 | 43.8 | 50.0 |
| | breeder populations | 8.0 | 80.0 | 12.0 |
| | Buckfast | 6.9 | 65.3 | 27.8 |
| | various breeds & hybrids | 15.3 | 76.4 | 8.3 |
| | mellifera | 45.8 | 45.8 | 8.3 |
| | localBreed | 35.6 | 57.8 | 6.7 |
| WinterDiapause | jellyBreed | 26.3 | 47.4 | 26.3 |
| | carnica | 6.2 | 50.0 | 43.8 |
| | breeder populations | 4.0 | 84.0 | 12.0 |
| | Buckfast | 18.1 | 59.7 | 22.2 |
| | various breeds & hybrids | 19.4 | 66.7 | 13.9 |
| | mellifera | 37.5 | 54.2 | 8.3 |
| | localBreed | 31.1 | 57.8 | 11.1 |

**Supplementary Table S4** Importance rating frequency per beekeeping profile variable with a significant effect for various traits

| Trait | Profile variable | | Importance rating Useless | Useful | Essential |
|---|---|---|---|---|---|
| Honey yield | Organic certification | Yes | 4.5 | 45.9 | 49.5 |
| | | No | 6.2 | 38.9 | 54.9 |
| Fecundity | | Yes | 22.5 | 66.7 | 10.8 |
| | | No | 14.8 | 60.5 | 24.7 |
| Calmness | | Yes | 18.9 | 75.7 | 5.4 |
| | | No | 16.7 | 60.5 | 22.8 |
| Disease resistance | Farm productions | Honey only | 8.8 | 61.4 | 29.8 |
| | | Honey, and other hive product(s) | 7.9 | 47.4 | 44.7 |
| | | Honey & pollination, & other hive product(s) | 0 | 50 | 50 |
| | | Honey & queens or swarms | 10.6 | 59.6 | 29.8 |
| | | Honey & queens or swarms, & other hive product(s) | 2.5 | 37.5 | 60 |
| | | Honey & queens or swarms & pollinatrion, & other hive product(s) | 2.9 | 64.7 | 32.4 |
| | | Honey & royal jelly, & other hive product(s) | 6.2 | 81.2 | 12.5 |
| | | Honey & royal jelly & queens or swarms, & other hive product(s) | 0 | 48 | 52 |
| Swarling tendency | Experience | Junior | 10 | 38.6 | 51.4 |
| | | Intermediate | 6 | 59.7 | 34.3 |
| | | Proficient | 5.9 | 70.6 | 23.5 |
| | | Master | 12.2 | 41.5 | 46.3 |
| | | Senior | 9.1 | 54.5 | 36.4 |
| Honey yield | Level of professionalizatin | hobbyist | 10 | 62 | 28 |
| | | small semi-professional | 13.6 | 40.9 | 45.5 |
| | | large semi-professional | 0 | 44.4 | 55.6 |
| | | small professional | 4.5 | 45.5 | 50 |
| | | medium professional | 4.4 | 36.8 | 58.8 |
| | | large professional | 3.2 | 27.4 | 69.4 |
| Winter feed autonomy | | hobbyist | 28 | 62 | 10 |
| | | small semi-professional | 9.1 | 68.2 | 22.7 |
| | | large semi-professional | 7.4 | 70.4 | 22.2 |
| | | small professional | 4.5 | 63.6 | 31.8 |
| | | medium professional | 10.3 | 58.8 | 30.9 |
| | | large professional | 12.9 | 64.5 | 22.6 |

Categories were as follows: junior (0 to 4 years), intermediate (5 to 9 years), proficient (10 to 14 years), master (15 to 24 years), senior (25 years and more); hobbyist (no revenue), small semi-professional (1 to 100 colonies), large semi-professional (101 to 280), small professional (1 to 200 colonies), medium professional (201 to 400 colonies), large professional (401 to 1800).

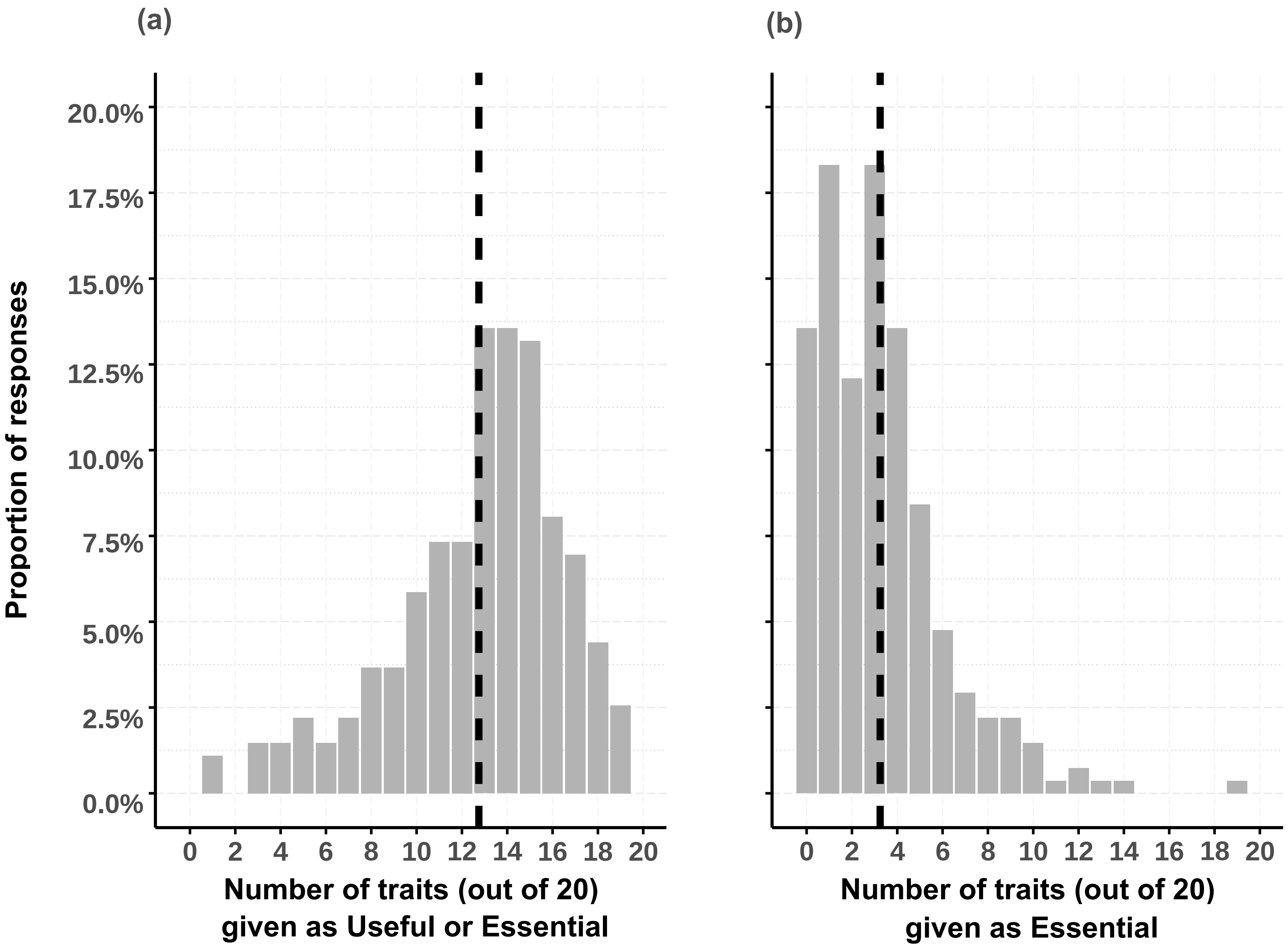


**Supplementary figure S1** Distribution of the number of traits beekeepers considered useful or essential (a) or essential only (b) to be included in the breeding goal.

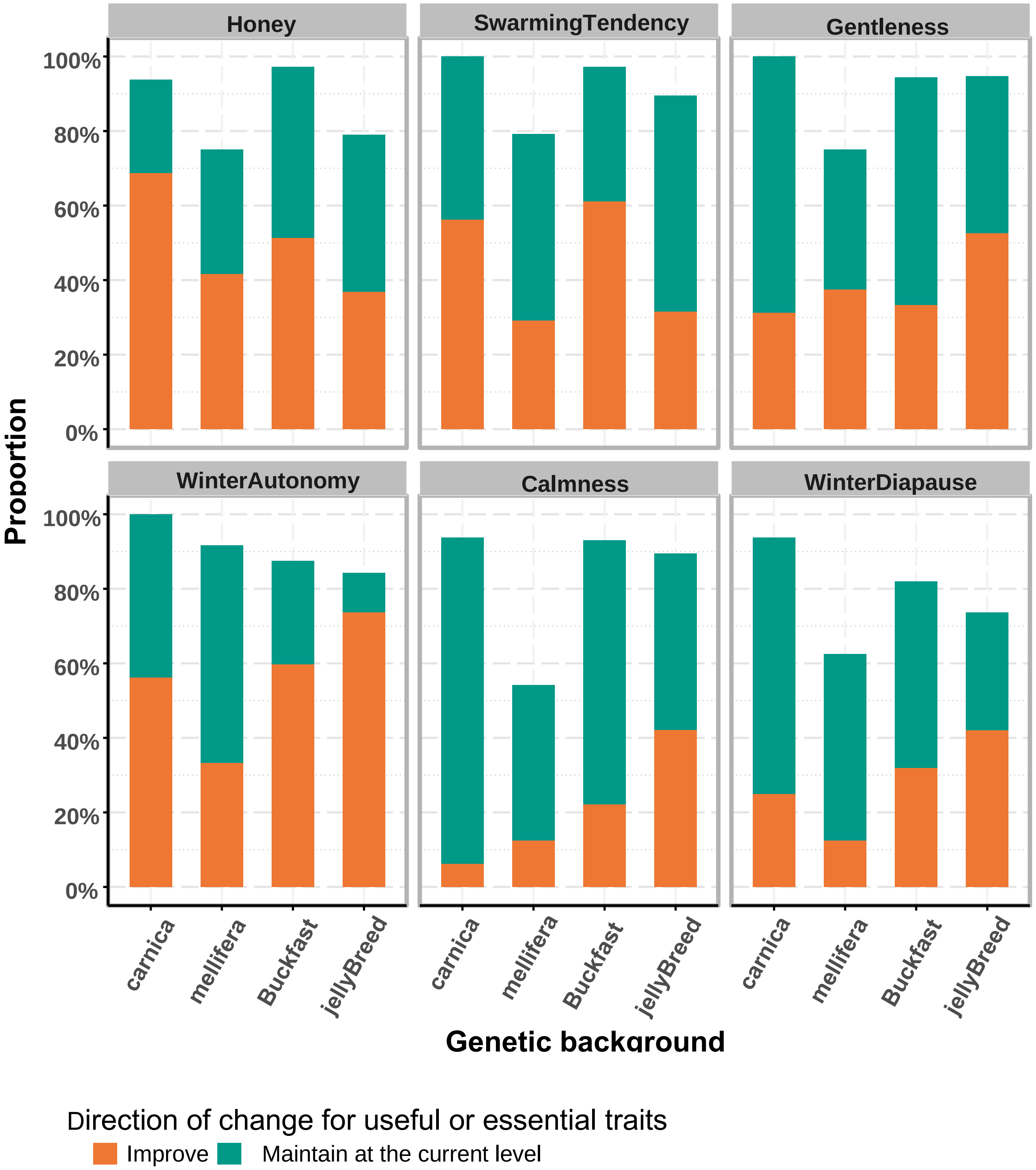


**Supplementary figure S2** Respondent proportion of desired trait genetic change for traits whose importance is influenced by bees' genetic background. Honey: honey yield; SwarmingTendency: swarming tendency; Gentlness: propensity of bees to not attack beekeepers during colony visit; WinterAutonomy: winter feed autonomy (no need of feeding); Calmness: propensity of bees to remain calm on the frames during colony visit, and not to run and cluster together; WinterDiapause: winter diapause (reduced colony activity, including egg-lay reduction or absence).
